# On Monitoring Brain Health from the Depths of Sleep: Feature Engineering and Machine Learning Insights for Digital Biomarker Development

**DOI:** 10.1101/2024.02.27.581950

**Authors:** Brice V McConnell, Yaning Liu, Ashis K Biswas, Brianne M. Bettcher, Lindsey M. Medenblik, Josiane L Broussard, Brendan P. Lucey, Alberto R. Ramos, Vitaly O. Kheyfets

## Abstract

**Backgroun:** Single-channel sleep electroencephalography (EEG) is a promising technology for creating cost-effective and widely accessible digital biomarkers for monitoring brain health. Sleep, notable for its numerous connections to brain health, is of particular interest in this context. Indeed, several of the best studied and widely recognized risk factors for neurodegenerative disease are also connected to aspects of sleep physiology, including biological sex, hypertension, diabetes, obesity/metabolic dysregulation, and immune system dysfunction. In this study, we utilize the unique signal characteristics of slow wave sleep (SWS) oscillatory events as features in machine learning models to predict underlying biological processes that are highly relevant to brain health. Our objective is to establish a foundation for algorithms capable of effectively monitoring physiological processes in sleep that directly and indirectly inform brain health using single-channel sleep EEG as a functional metric of brain activity.

**Methods:** Utilizing data from the Cleveland Family Study, we analyzed 726 overnight polysomnography recordings to extract features from slow waves and adjacent oscillatory events. Advanced signal processing and machine learning techniques, including random forest models, were employed to engineer features and predict health-related outcomes such as age, cerebrovascular risk factors, endocrine functions, immune system activity, and sleep apnea.

**Results:** Our models demonstrated significant predictive capability for several outcomes, including age (R^2^ = 0.643, p < 0.001), and sex classification (area under the receiver operator characteristic (AUROC) curve = 0.808), diabetes and hypertension diagnosis (AUROC = 0.832 and 0.755, respectively). Significant predictions were also modeled for metabolic/endocrine functions (including blood concentrations of IGF-1, leptin, ghrelin, adiponectin, and glucose), and immune markers (including IL-6, TNF-alpha, and CRP). In addition, this approach provided successful predictions in regression modeling of BMI and both regression and classification of sleep apnea.

**Discussion:** This study demonstrates the potential of using features from oscillatory events in single-channel sleep EEG as digital biomarkers. These biomarkers can identify key health and demographic factors that both affect brain health and are indicative of core brain functions. By capturing the complex interactions of neural, metabolic, endocrine, and immune systems during sleep, our findings support the development of single-channel EEG as a practical tool for monitoring complex biological processes through metrics that originate in brain physiology. Future research should aim to refine these digital biomarkers for broader home-based applications that may utilize inexpensive “wearable” devices to provide a scalable and accessible tool for tracking brain health-related outcomes.

## Introduction

Electroencephalography (EEG) is a core method in neurophysiology and clinical neuroscience, capturing the brain’s electrical activity through scalp electrodes, and providing a non-invasive method to assess cerebral activities underlying brain communication^1^. EEG is increasingly recognized as a viable digital biomarker for monitoring brain health and detecting neurological disease^2,3^. While there are promising applications for this technology during wake-state EEG, its application during wakefulness encounters significant challenges due to the complexity of brain processes and the mixed signals they produce, which imposes significant limits on the development of single-channel EEG biomarkers^4^. These limitations create formidable barriers to practical applications for the technology, where arrays of multiple recording electrodes are impractical in the self-application of EEG devices that might be used for long-term brain health monitoring. In contrast to wake EEG, sleep EEG exhibits several characteristics that expand the ability to measure more focused aspects of brain communication from a single channel of EEG, including more synchronous and predictable oscillatory patterns^5^. Thus, sleep EEG provides a unique set of signal qualities that are suitable to build digital markers using simple EEG recording equipment, such as a single channel headband “wearable” device.

Sleep EEG is defined by distinctive patterns that reflect subcortical circuit coordination and large-scale synchronous cortical activities^6–8^. Within the brain, sleep contributes to synaptic regulation and memory consolidation^9,10^. Additionally, the neurophysiology of sleep has numerous connections to other physiological systems, including metabolic regulation^11^, endocrine functions^12,13^ and the immune system^14,15^. Notably, the interconnected relationships between the functions of sleep in the brain and in the body’s peripheral systems are crucial for protecting brain health. Conditions such as obesity, diabetes, and hypertension, which can be driven by sleep disruption^16–18^, are also significant risk factors for neurodegenerative diseases, including Alzheimer’s disease and vascular dementia^19,20^. Further, the regulation of immune functions during sleep highlights the complex interplay between sleep, systemic inflammation, and risk of neurodegenerative disease^21^. These properties of sleep provide a rationale and mechanistic framework for the use of sleep EEG’s oscillatory signals as a digital biomarker to inform core biological processes that impact brain health. By monitoring variations in brain activity during sleep, single channel sleep EEG offers a unique window into the systemic and neurophysiological underpinnings of brain health, presenting a promising avenue for early detection and intervention in conditions predisposing individuals to neurodegenerative diseases. Within sleep, slow-wave sleep (SWS), in particular, plays a critical role in the central nervous system (CNS) and other bodily functions^8^. Indeed, the neuroprotective properties of SWS make it a target for interventions designed to prevent and treat neurodegenerative disease^22^. SWS is comprised of discrete oscillatory events representing bursts of activity in cortical and subcortical brain regions^6,23^. Several types of these oscillatory events have been described, including slow waves, spindles, theta bursts, and ripples^6,23–26^. The morphology of oscillatory event waveforms, and the temporal coupling of these events to one-another, demonstrate distinct patterns with aging and neurodegenerative disease^27–32^. There are also well-characterized oscillatory event properties that occur in association with biological processes and are highly related to SWS physiology, including memory performance^33^, sex differences^34–38^, and insulin sensitivity^11,39,40^. Oscillatory events also have distinct anatomical correlations that inform underlying brain circuits and provide specificity to the origins of EEG oscillations^24,25,41–44^. Thus, incorporating oscillatory event-based features into predictive models provides a foundation for understanding the neurophysiological mechanisms that support SWS-based digital biomarkers in brain health monitoring.

While there are numerous descriptions of oscillatory event morphology and frequency/power properties correlating with underlying brain physiology, it is unclear how these observations may inform the construction of a digital biomarker to predict brain health-related outcomes. Here we capitalized on domain knowledge to engineer features from slow waves and adjacent oscillatory events, including theta bursts and spindles, from single-channel sleep EEG to construct components of a digital biomarker. Statistical metrics of waveform properties, time-frequency relationships, and EEG power analyses were converted into feature sets from each overnight recording. These features were then applied within random forest machine learning models to predict outcomes known to be related to SWS and relevant to monitoring brain health. We tested the predictive performance in several sleep-related outcomes (including age, cerebrovascular risk factors, endocrine functions, and immune system activity) to examine the ability of these oscillatory event-based features to capture elements of complex physiological constructs that are highly relevant to applications of brain health monitoring.

## Methods

### Participant data

Participant data from the Cleveland Family Study (CFS) was accessed through the National Sleep Research Resource^45,46^. All data were collected with written informed consent under protocols approved by a local institutional review board. Single-channel EEG data within overnight polysomnography recordings were utilized from visit 5 study data. A total of 735 records were available, although 9 were excluded from analysis due to data quality issues in recordings and missing EEG data. The remaining 726 recordings were used for model training and testing, although not all recordings were associated with valid participant data for each variable. Data entries for variables that were beyond the assay floor or ceiling limitations of instrument measurement were excluded from analyses, as were values of zero in data sets requiring logarithmic conversion. To minimize the potential impact of extreme data points that may skew the regression modeling and affect the model’s predictive accuracy and generalizability, we also eliminated outlier variable data points that fell below 1.5 times the interquartile range (IQR) below the 25th percentile or above 1.5 times the IQR above the 75th percentile.

The remaining data was divided into sets for training/validation of the models and independent testing of the model performance. The training/validation sets were used exclusively for training the models and optimizing hyperparameters, while the testing sets were used to test predictive performance of the models using segregated data that was excluded from model training. Separately, for each valid outcome variable, we randomly assigned participant family IDs into training/validation and testing sets in an approximately 80/20 ratio, respectively. This assignment based on family IDs ensured the complete segregation of family IDs into either the training/validation or testing set. This strategy was implemented due to the large number of directly related participants in the cohort to: 1) prevent leakage of familial traits that might impact the model performance in the testing sets, and 2) create a distinct training and testing set for each model for improved generalizability of our findings.

Details of cohort recruitment and study design, including molecular assays, body measurements, and health history acquisition, are previously described for the CFS cohort, visit 5^45,47,48^. Distributions of prediction variables were visually inspected, and variables with highly skewed distributions were converted to their natural log values for predictive regression. Sleep apnea as assessed by apnea hypopnea index (AHI), defined by all apneas and hypopneas with > 30% flow reduction and >= 3% oxygen desaturation and with or without arousal per hour of sleep, in accordance with guidelines from the American Academy of Sleep Medicine (AASM)^49^. Classification of apnea severity used established threshold values of at least 5 AHI for mild category, at least 15 AHI for moderate, and at least 30 AHI for severe^49^.

### EEG data acquisition

Polysomnography and sleep staging for CFS are previously described^47^. Briefly, signal acquisition was performed using a 14-channel Compumedics E-Series System (Abbotsford, Australia). EEG was obtained using gold cup electrodes at a sampling rate of 128 Hz (filtered high pass 0.16 Hz and low pass 64 Hz), and electrode pair C3-A2 was used for all analysis (prior analyses of oscillatory events demonstrate similar oscillatory event composition between C3-A2 and C4-A1^50^).

### Slow wave event identification

All signal processing steps were performed within MATLAB R2021b (MathWorks, Inc., Natick, MA). SWs were identified via automated zero-crossing detection as previously described^32,42,50^. Briefly, SW detection was performed from C3-A2 electrode pair in sleep stages N2 and N3. Automated management of high amplitude artifacts was accomplished via exclusion of EEG segments exceeding 900 μV after detrending data with sliding window of three seconds across raw data. Next, EEG data were detrended and band-pass filtered in a forward and backward direction using a sixth-order Butterworth filter between 0.16 and 4 Hz. Zero crossings were identified to detect negative and positive half-waves, and SW events were identified when the half-wave pairs approximated a frequency range of 0.4–4 Hz. Minimal and maximal half-wave amplitudes were measured, and SWs with both positive and negative maximum amplitudes in the top 50% of all waves were selected for subsequent coupling analysis, providing a dynamic threshold for event identification. An upper threshold of ±200 μV for zero crossing pairs was utilized to reduce misidentification of non-SW events. A further reduction of false identifications was accomplished by rejecting all zero crossing pairs with peak/trough amplitudes exceeding four standard deviations from the mean min/max zero crossing pair values for each subject.

### Time-frequency (TF) spectrogram generation

Time-frequency (TF) wavelet spectrograms of SW-associated time windows were created via established methods^32,50^. Briefly, the troughs of each SW were centered in 5-s windows of EEG data and matched baseline intervals. A Morlet-wavelet transformation (65 cycles from 4 to 10 Hz) was applied to the unfiltered EEG for SW and baseline segments between 4 and 20 Hz in steps of 0.25 Hz with varying wave numbers (65 cycles from 4 to 10 Hz with a step size of 0.0938 to match the frequency step size). The mean of the baseline windows was used to normalize the power of the mean Morlet-wavelet SW windows.

### Slow wave selection by thresholding

We developed a computational approach to analyze EEG slow waves using bandpass filtering, Empirical Mode Decomposition (EMD), and Short-Time Fourier Transform (STFT) functions. Initially, EEG time series data were bandpass filtered via the MATLAB bandpass function using a finite impulse response (FIR) filter within pre-defined frequency ranges, separately for time windows immediately before (PreSW: 4.5-8 Hz and 8-11 Hz) and after (PostSW: 10-14 Hz, 12-16 Hz, and 14-18 Hz) the trough of the slow waves. The bandpass filtered data were subjected to EMD to extract the first Intrinsic Mode Function (IMF) of each time window, which represents the dominant oscillatory mode. Subsequently, we computed the STFT values for these IMFs in specific time segments relative to the trough of the slow waves. For the PreSW period, STFT was calculated from 0.5 seconds before to the trough (time zero), and for the PostSW period, STFT was calculated from the trough to 0.5 seconds and 0.5 to 1 second after the trough. To focus on slow waves with significant spectral content, we implemented a thresholding procedure. We identified and retained slow waves with the highest 25% of STFT values, which represent periods with high spectral power in the specified frequency bands and time regions. This subset of slow waves was then used for further analysis.

### Slow wave clustering by unsupervised machine learning

An unsupervised machine learning approach was implemented to reduce heterogeneity of slow wave feature sets by clustering slow waves using feature characteristics. First, the bandpass MATLAB function with FIR was applied to the time series data using eight distinct frequency ranges: 0.5-1.5 Hz, 0.8-1.8 Hz, 1.1-2.1 Hz, 1.4-2.4 Hz, 1.7-2.7 Hz, 2-3 Hz, 2.3-3.3 Hz, and 2.7-3.7 Hz. This provided eight different bandpass instances for each time series window to utilize for slow wave feature extraction. For each of these bandpassed time series windows, the first IMF was computed using EMD, resulting in a set of concatenated IMFs. After the IMF extraction, waveform features were derived from these concatenated IMFs. Waveform event markers included the identification of the closest trough to the middle point of the window, peaks on either side of this trough, zero crossings around these peaks and the trough, and the distances between these zero crossings. Additionally, amplitudes at the identified peaks and trough, as well as slopes between these points, were calculated.

Outlier slow wave events, defined as containing waveform values exceeding seven standard deviations from the mean, were excluded from further analysis. The spectrogram data, resized to 10×10 dimensions for each window, underwent a similar outlier detection process, with outlier slow waves defined as containing spectrogram values deviating more than eight standard deviations from the mean in normalized EEG power. Normalization was independently applied to both waveform features and STFT values. This was achieved by calculating the mean and standard deviation for each feature type and then applying z-score normalization. The normalized features from the waveform and STFT values were then combined into a single feature set for each time-series window.

The combined features from all processed files were used in a k-means clustering algorithm to categorize the slow waves into two clusters. The number of clusters was predefined as two for this analysis. A universal set of reference centroid characteristics was created to ensure consistent k-means cluster assignment across data files. These reference centroids, derived from the pooled data of ten recordings, were subsequently employed to categorize slow waves in each individual recording of the entire study dataset. This uniform cluster assignment ensured a consistent analytical approach across subsequent processing steps with cluster 1 and cluster 2 slow wave events. This clustering divided the slow waves for each participant into two clusters for subsequent feature extraction, reducing the heterogeneity of slow wave features within each of the subsequent feature extraction steps.

### Feature extraction from slow wave waveforms

Waveform metrics from slow waves in both cluster 1 and cluster 2 of the k-means clustering process were subsequently processed to create statistical summary features. The waveform metrics utilized for these features included distances between zero crossings, amplitudes at peaks and troughs, slopes, and positions of peaks, troughs, and zero crossings. For each cluster of slow waves, these waveform metrics were further categorized based on their frequency characteristics into low (0.5-1.5 Hz, 0.8-1.8 Hz), mid (1.1-2.1 Hz, 1.4-2.4 Hz, 1.7-2.7 Hz), and high (2-3 Hz, 2.3-3.3 Hz, 2.7-3.7 Hz) frequency feature groups. Statistical analyses were then conducted on these grouped metrics to extract thirteen statistical measures for each metric group: mean, standard deviation, skewness, kurtosis, range, median, mode, variance, mean absolute deviation (MAD), interquartile range, 25th percentile, 75th percentile, and root mean square (RMS). These statistical tests were computed separately for slow waves in cluster 1 and cluster 2 from k-means, resulting in 1,170 statistical summary features.

### Feature extraction from slow wave TF spectrograms

We processed individual spectrograms through a series of image processing steps in the MATLAB image processing toolbox to obtain image-based feature metrics as previously described^32^. Briefly, each spectrogram, resized to a 100×100 pixel image, was converted into a binary image. Next, boundaries of oscillatory events were identified in the spectrogram images after removing noise, gaps, and holes. From these boundaries of oscillatory events, we computed region properties, including centroid, area, perimeter, major and minor axis lengths, and pixel values within the detected regions.

We next defined five time-frequency (TF) windows based on specific time-frequency coordinates. The first TF-window spanned the preSW time from -1.4 to -0.3 seconds relative to the SW trough and covered a frequency range from 11 Hz to 18 Hz. The second TF-window was positioned around the SW trough, extending from -0.6 seconds to 0.3 seconds, with its frequency range set between 4 Hz and 8 Hz. The third TF-window, covering the same time span as the second, encompassed frequencies from 8 Hz to 11 Hz. The fourth TF-window ranged from 0.3 to 1.4 seconds in time, between 9 Hz and 13 Hz in frequency. The fifth TF-window, also postSW, extended from 0.3 to 1.4 seconds in time but captured higher frequencies, from 13 Hz to 18 Hz. We calculated ratios and percentages of events occurring within and across these regions, offering insights into the distribution and interaction of events in different parts of the spectrogram. Additionally, we assessed centroid standard deviations and average inter-centroid distances within each region, adding another layer of spatial analysis.

The processed data from each spectrogram also underwent a comprehensive statistical analysis. This included the extraction of advanced features such as event density, average inter-event distance, average event duration, frequency spread, temporal dynamics, roundness, and average pixel intensity of the detected events. We calculated event density by summing the areas of all detected events and normalizing this by the total image size. The average event duration and frequency spread were approximated using the major and minor axis lengths of the detected events. Temporal dynamics were assessed by analyzing the differences in the centroids of consecutive events. Roundness was computed as a measure of event shape variability, and the average pixel intensity provided insight into the event intensity profile. A nearest neighbor analysis also determined the average distance between each event and its closest neighboring event, providing a metric of spatial distribution. These metrics were computed separately for slow waves in clusters 1 and 2 from k-means, resulting in 72 features.

### Feature extraction from slow wave STFT metrics

We defined a set of seven bandpass frequency ranges: 4-6 Hz, 6-8 Hz, 8-10 Hz, 10-12 Hz, 12-14 Hz, 14-16 Hz, and 16-18 Hz to capture a broad spectrum of EEG activity near slow wave events. We next identified six distinct time regions relative to the preSW/postSW trough index point: - 1.5 to -1.0 seconds, -1.0 to -0.5 seconds, -0.5 seconds to trough index, trough index to +0.5 seconds, +0.5 to +1.0 seconds, and +1.0 to +1.5 seconds. The data within each region underwent STFT analysis, and these values were used to compute four statistical measures for the STFT data in each band and region: mean, standard deviation, skewness, and kurtosis. These metrics were computed separately for slow waves in clusters 1 and 2 from k-means, resulting in 336 features.

### Random forest machine learning classification and regression

We created Random Forest (RF) regression and classification models using the TreeBagger function within the Statistics and Machine Learning Toolbox of MATLAB (R2021b). A total of 1,578 features extracted each EEG recording were used as input to the models. We split the data into training and validation sets, using 80% of the available training/validation data for training and the remaining 20% for validation of hyperparameter settings (see Tables 1 and 2). For RF regression and classification, we applied forty cycles of Bayesian optimization to optimize the hyperparameters, including the number of trees, the number of predictors to sample, maximum number of splits, minimum leaf size, and in-bag fraction. For RF classification, class weights were optimized as additional parameters within the TreeBagger function during training. Upon identifying the best hyperparameters, we trained each RF model using the optimized settings on the entire feature matrix, encompassing both the training and validation sets, to fully leverage the available data. The performance of each trained model was then evaluated using separate test sets of participants, and all values reported for model performance were obtained in these tests. For RF regression, metrics including R^2^, p-values, mean absolute error (MAE), mean squared error (MSE), root mean squared error (RMSE) and the mean absolute percentage error (MAPE) were computed. For RF classification, we computed area under the receiver operating characteristic (AUROC) curve, accuracy, precision, recall and F1 score.

**Table 1.** Subject Selection.

| Variable | #<br>excluded<br>for lack<br>of data | Total<br>*N*<br>used | *N*<br>training<br>set | Median age (IQR)<br>training set | *N* (%)<br>female<br>training<br>set | *N*<br>test<br>set | Median age (IQR)<br>test set | *N* (%)<br>female<br>test set |
| --- | --- | --- | --- | --- | --- | --- | --- | --- |
| Apnea | 77 | 658 | 510 | 42.819<br>(21.851 – 53.107) | 287<br>(56.28) | 148 | 43.964<br>(22.467 – 57.156) | 93<br>(62.84) |
| ln-Apnea | 88 | 647 | 502 | 42.97<br>(22.234 – 53.214) | 280<br>(55.78) | 145 | 44.512<br>(23.261 – 57.158) | 90<br>(62.07) |
| Biological<br>Brain Age | 9 | 726 | 602 | 43.182<br>(23.55 – 54.96) | 338<br>(56.15) | 124 | 46.88<br>(24.84 – 54.98) | 62<br>(50) |
| Biological<br>Sex | 9 | 726 | 568 | 43.127<br>(22.455 – 54.343) | 305<br>(53.70) | 158 | 45.491<br>(25.615 – 58.066) | 95<br>(60.13) |
| BMI | 21 | 714 | 600 | 43.561<br>(23.769 – 54.999) | 330<br>(55) | 114 | 43.153<br>(22.130 – 56.652) | 62<br>(54.39) |
| Diabetes | 12 | 723 | 573 | 44.841<br>(25.503 – 56.016) | 307<br>(53.58) | 150 | 37.986<br>(18.844 – 51.818) | 91<br>(60.67) |
| Fasting<br>Glucose | 96 | 639 | 515 | 41.812<br>(21.796 – 53.526) | 293<br>(56.89) | 124 | 42.914<br>(24.602 – 54.667) | 65<br>(52.42) |
| Fasting<br>Insulin | 71 | 664 | 540 | 43.183<br>(23.611 – 55.219) | 296<br>(54.81) | 124 | 45.851<br>(24.322 – 55.104) | 73<br>(58.87) |
| Ghrelin | 49 | 686 | 545 | 43.149<br>(22.491 – 54.798) | 298<br>(54.68) | 141 | 43.647<br>(24.131 – 54.634) | 74<br>(52.48) |
| Hypertension | 17 | 718 | 520 | 43.967<br>(24.277 – 54.673) | 289<br>(55.58) | 198 | 42.324<br>(21.307 – 55.277) | 107<br>(54.04) |
| ln-adiponectin | 48 | 687 | 527 | 43.036<br>(23.797 – 54.623) | 283<br>(53.70) | 160 | 44.702<br>(24.387 – 55.123) | 91<br>(56.88) |
| ln-CRP-am | 76 | 659 | 503 | 46.459<br>(29.359 – 56.048) | 281<br>(55.86) | 156 | 39.985<br>(19.121 – 53.435) | 89<br>(57.05) |
| ln-IGF-1am | 30 | 705 | 564 | 42.930<br>(23.986 – 54.520) | 299<br>(53.01) | 141 | 45.678<br>(22.026 – 57.955) | 88<br>(62.41) |
| ln-IL-1b | 151 | 584 | 481 | 42.817<br>(22.253 – 53.966) | 269<br>(55.83) | 103 | 44.527<br>(24.730 – 56.207) | 61<br>(59.22) |
| ln-IL-6am | 44 | 691 | 502 | 42.995<br>(22.981 – 54.366) | 267<br>(53.19) | 189 | 44.802<br>(24.326 – 55.299) | 108<br>(57.14) |
| ln-IL-10am | 101 | 634 | 493 | 43.036<br>(22.491 – 54.724) | 265<br>(53.75) | 141 | 43.948<br>(23.307 – 55.099) | 82<br>(58.16) |
| ln-Leptin-am | 61 | 674 | 495 | 44.465<br>(23.926 – 55.020) | 265<br>(53.54) | 179 | 43.055<br>(23.288 – 54.783) | 105<br>(58.66) |
| ln-TGFalpha | 67 | 668 | 567 | 43.313<br>(22.684 – 54.825) | 312<br>(55.03) | 101 | 44.841<br>(24.652 – 57.155) | 54<br>(53.47) |
| Oral GTT | 177 | 558 | 415 | 40.810<br>(22.435 – 52.240) | 231<br>(55.66) | 143 | 40.906<br>(19.514 – 52.430) | 84<br>(58.74) |

**Table 2.** Classification Category Characteristics.

| Prediction Variable | Training *N*<br>Category 1 | Training *N*<br>Category 2 | Test set *N*<br>Category 1 | Test set *N*<br>Category 2 |
| --- | --- | --- | --- | --- |
| Biological Sex | 263 | 305 | 63 | 95 |
| Diabetes | 85 | 488 | 14 | 136 |
| Hypertension | 166 | 354 | 148 | 50 |
| Mild Apnea | 229 | 281 | 76 | 72 |
| Moderate Apnea | 95 | 415 | 33 | 115 |
| Severe Apnea | 17 | 493 | 6 | 142 |

*Biological Sex: Category 1 = male, Category 2 = female; Diabetes: Category 1 = diagnosed with diabetes, Category 2 = no diagnosis of diabetes; Hypertension: Category 1 = diagnosed with hypertension, Category 2 = no diagnosis of hypertension; Mild Apnea: Category 1 = AHI equal to or greater than mild apnea, Category 2 = AHI less than mild apnea; Moderate Apnea: Category 1 = AHI equal to or greater than moderate apnea, Category 2 = AHI less than moderate apnea; Severe Apnea: Category 1 = AHI equal to or greater than severe apnea, Category 2 = AHI less than or equal to severe apnea.*

## Results

### Age Predictive Modeling

Random forest regression modeling, when applied to predict age in an independent test set of participants, achieved an R² of 0.643 (p < 0.001), indicating high predictive performance in modeling variance of age (Figure 1; Table 3). The model displayed a mean absolute error (MAE) of 8.651, a mean squared error (MSE) of 122.098, a root mean squared error (RMSE) of 11.050, and a mean absolute percentage error (MAPE) of 25.031% (Table 3).

**Figure 1.**
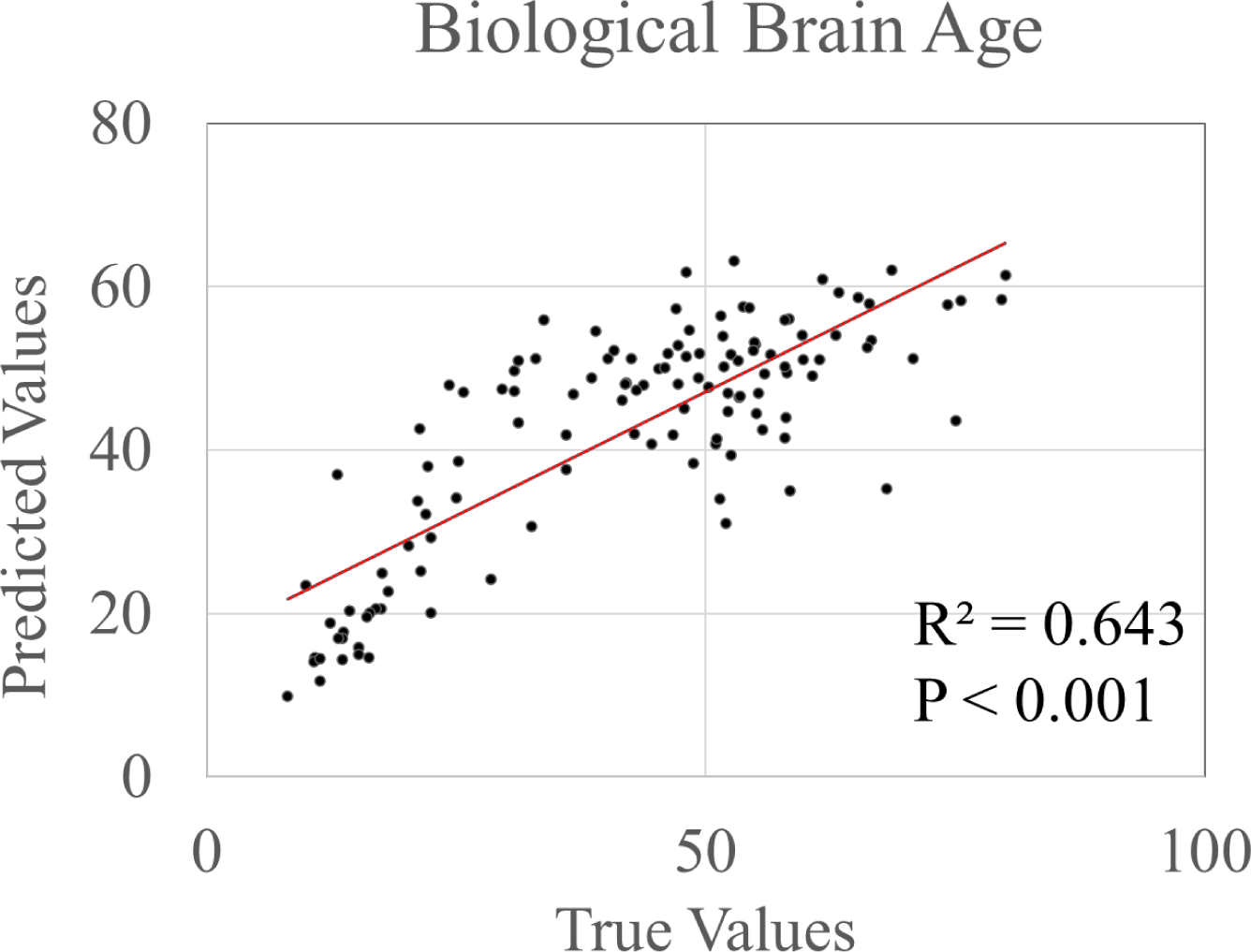
Regression plot of the Biological Brain Age predictive test.

**Table 3.** Regression Results.

| Test | R square | Significance F | MAE | MSE | RMSE | MAPE |
| --- | --- | --- | --- | --- | --- | --- |
| Apnea | 0.240 | p < 0.000 | 0.924 | 1.357 | 1.165 | 129.173% |
| Biological Brain Age | 0.643 | p < 0.001 | 8.651 | 122.098 | 11.050 | 25.031% |
| BMI | 0.200 | p < 0.001 | 6.354 | 60.095 | 7.752 | 22.868% |
| Fasting Glucose | 0.162 | p < 0.001 | 6.734 | 78.292 | 8.848 | 7.109% |
| Fasting Insulin | 0.027 | p = 0.070 | 0.437 | 0.273 | 0.523 | 19.991% |
| Ghrelin | 0.075 | p = 0.001 | 282.297 | 128018.144 | 357.796 | 33.272% |
| ln-adiponectin | 0.078 | p < 0.001 | 0.498 | 0.370 | 0.609 | 10.493% |
| ln-CRP-am | 0.084 | p < 0.001 | 1.014 | 1.547 | 1.207 | 218.306% |
| ln-IGF-1am | 0.380 | p < 0.001 | 0.317 | 0.159 | 0.399 | 5.808% |
| ln-IL-1b | 0.006 | p = 0.436 | 0.714 | 0.877 | 0.936 | 127.201% |
| ln-IL-6am | 0.120 | p < 0.001 | 0.567 | 0.501 | 0.708 | 186.430% |
| ln-IL-10am | 0.007 | p = 0.338 | 0.861 | 1.187 | 1.089 | 375.164% |
| ln-leptin-am | 0.057 | p = 0.001 | 0.700 | 0.768 | 0.876 | 30.306% |
| ln-TGF-alpha | 0.054 | p = 0.020 | 0.514 | 0.381 | 0.617 | 110.426% |
| Oral GTT | 0.115 | p < 0.001 | 28.264 | 1323.087 | 36.374 | 25.152% |

95% CI. MAE = Mean Absolute Error; MSE = Mean Squared Error, RMSE = Root Mean Squared Error, MAPE = Mean Absolute Percentage Error

### Cerebrovascular Risk Factor Modeling

Random forest classification modeling for diabetes diagnosis yielded an AUROC of 0.832 and an overall accuracy of 57.3% (Figure 2; Table 4). Precision for predicting diabetes was 0.929, while for no diagnosis, it was 0.537. Recall scores were 0.171 for individuals diagnosed with diabetes and 0.986 for those without, leading to F1 scores of 0.289 and 0.695, respectively. For hypertension diagnosis, classification modeling resulted in an AUROC of 0.755 and an accuracy of 69.7%. Precision was 0.660 for those diagnosed with hypertension and 0.696 for those without. Recall for individuals with hypertension was 0.423, compared to 0.858 for those without hypertension, with F1 scores of 0.516 and 0.768, respectively. A random forest regression model predicting blood fasting glucose levels demonstrated an R² of 0.162 (p < 0.001), with a mean absolute error (MAE) of 6.734, mean squared error (MSE) of 78.282, root mean squared error (RMSE) of 8.848, and mean absolute percentage error (MAPE) of 7.109%. An independent regression model for blood glucose levels in an oral glucose tolerance test (OGTT) showed an R² of 0.115 (p < 0.001), MAE of 28.264, MSE of 1323.087, RMSE of 36.374, and MAPE of 25.152%. Another model focused on predicting BMI metrics reported an R² of 0.200 (p < 0.001), with an MAE of 8.651, MSE of 122.098, RMSE of 11.050, and a MAPE of 25.031%. The fasting insulin regression model showed an R² of 0.027 (p = 0.07), with an MAE of 0.437, MSE of 0.273, RMSE of 0.523, and a MAPE of 19.991%.

**Figure 2.**
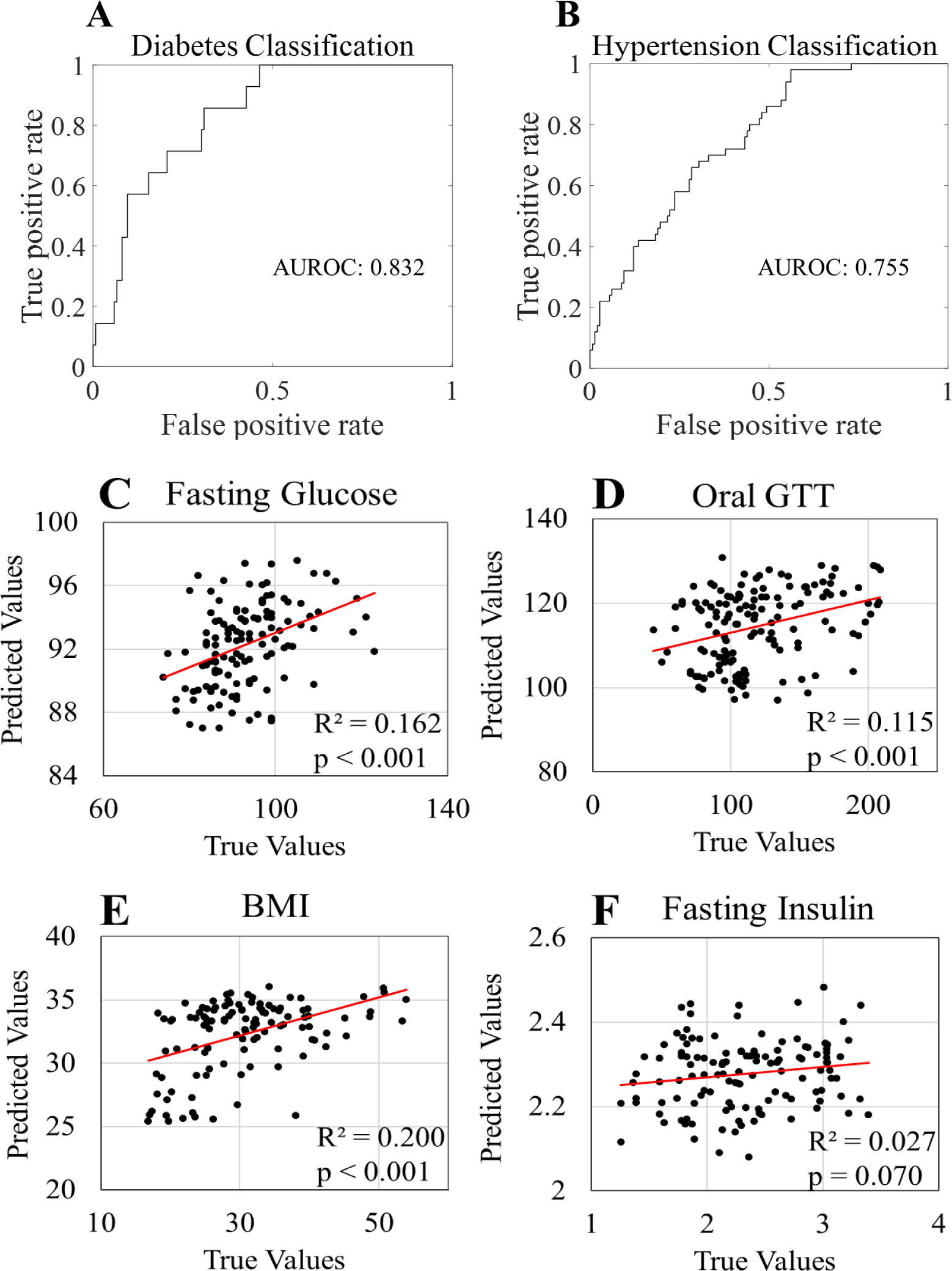
ROC plots for the (A) Diabetes and (B) Hypertension classification predictions, as well as the regression plots for the (C) Fasting Glucose, (D) Oral GTT, (E) BMI predictions, and (F) Fasting Insulin.

**Table 4.** ROC Metrics.

| Prediction Variable | Accuracy | AUROC | Precision Category 1 | Precision Category 2 | Recall 1 | Recall 2 | F1 Repeated 1 | F1 Repeated 2 |
| --- | --- | --- | --- | --- | --- | --- | --- | --- |
| Biological Sex | 0.766 | 0.808 | 0.779 | 0.746 | 0.822 | 0.691 | 0.80 | 0.718 |
| Diabetes | 0.573 | 0.832 | 0.929 | 0.537 | 0.171 | 0.986 | 0.289 | 0.695 |
| Hypertension | 0.697 | 0.755 | 0.660 | 0.696 | 0.423 | 0.858 | 0.516 | 0.768 |
| Mild Apnea | 0.689 | 0.764 | 0.868 | 0.5 | 0.647 | 0.783 | 0.742 | 0.610 |
| Moderate Apnea | 0.561 | 0.670 | 0.727 | 0.513 | 0.300 | 0.868 | 0.425 | 0.645 |
| Severe Apnea | 0.905 | 0.644 | 0.167 | 0.937 | 0.1 | 0.964 | 0.125 | 0.95 |

*Biological Sex: Category 1 = male, Category 2 = female; Diabetes: Category 1 = diagnosed with diabetes, Category 2 = no diagnosis of diabetes; Hypertension: Category 1 = diagnosed with hypertension, Category 2 = no diagnosis of hypertension; Mild Apnea: Category 1 = AHI equal to or greater than mild apnea, Category 2 = AHI less than mild apnea; Moderate Apnea: Category 1 = AHI equal to or greater than moderate apnea, Category 2 = AHI less than moderate apnea; Severe Apnea: Category 1 = AHI equal to or greater than severe apnea, Category 2 = AHI less than or equal to severe apnea.*

### Endocrine System Modeling

Random forest classification modeling accurately predicted biological sex, achieving an AUROC of 0.808 and an overall accuracy of 76.6% (Figure 3; Table 4). Precision values were 0.779 for predicting males and 0.746 for predicting females. Recall scores were reported as 0.822 for males and 0.691 for females. Consequently, F1 scores were calculated to be 0.800 for males and 0.718 for females, indicating the model’s balanced performance in predicting sex.

**Figure 3.**
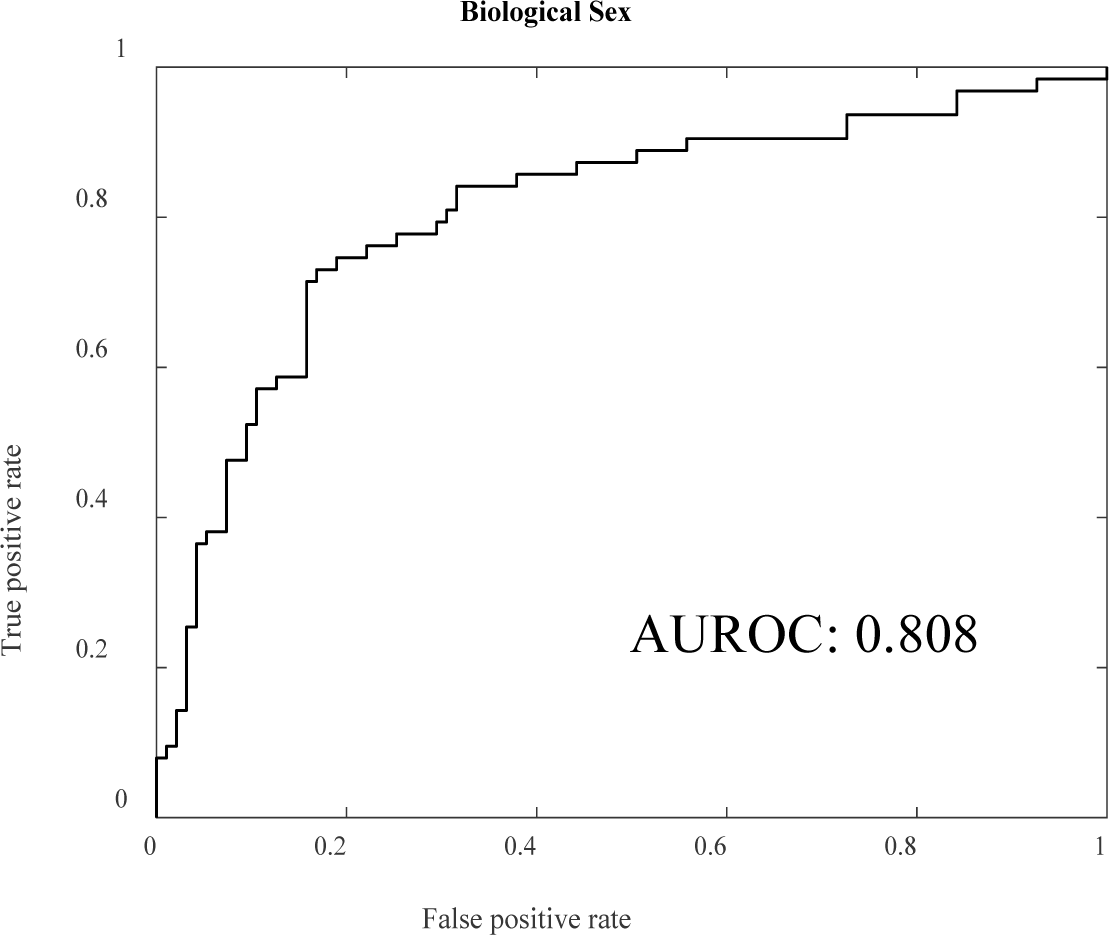
ROC plot for Biological Sex prediction.

Regression analysis for blood ln-IGF-1 levels achieved an R² of 0.380 (p < 0.001), suggesting a substantial portion of ln-IGF-1 level variability, approximately 38%, could be accounted for by the model (Figure 4; Table 3). The MAE for this prediction was 0.317, MSE was 0.159, RMSE was calculated as 0.399, and the MAPE was 5.808%. The model predicting blood adiponectin levels using a predictive random forest model yielded an R² of 0.0778 (p < 0.001), indicating the model explained approximately 7.78% of the variance in adiponectin levels. The mean absolute error (MAE) was reported at 0.498, with a mean squared error (MSE) of 0.370, root mean squared error (RMSE) of 0.609, and a mean absolute percentage error (MAPE) of 10.493%. Similar predictive performance was observed for blood ghrelin levels, with an R² of 0.075 (p = 0.001). The analysis revealed an MAE of 282.297, MSE of 128,018.144, RMSE of 357.796, and a MAPE of 33.272%. For blood leptin levels, the regression model demonstrated an R² of 0.057 (p = 0.001), with an MAE of 0.700, MSE of 0.768, RMSE of 0.876, and a MAPE of 30.306%.

**Figure 4.**
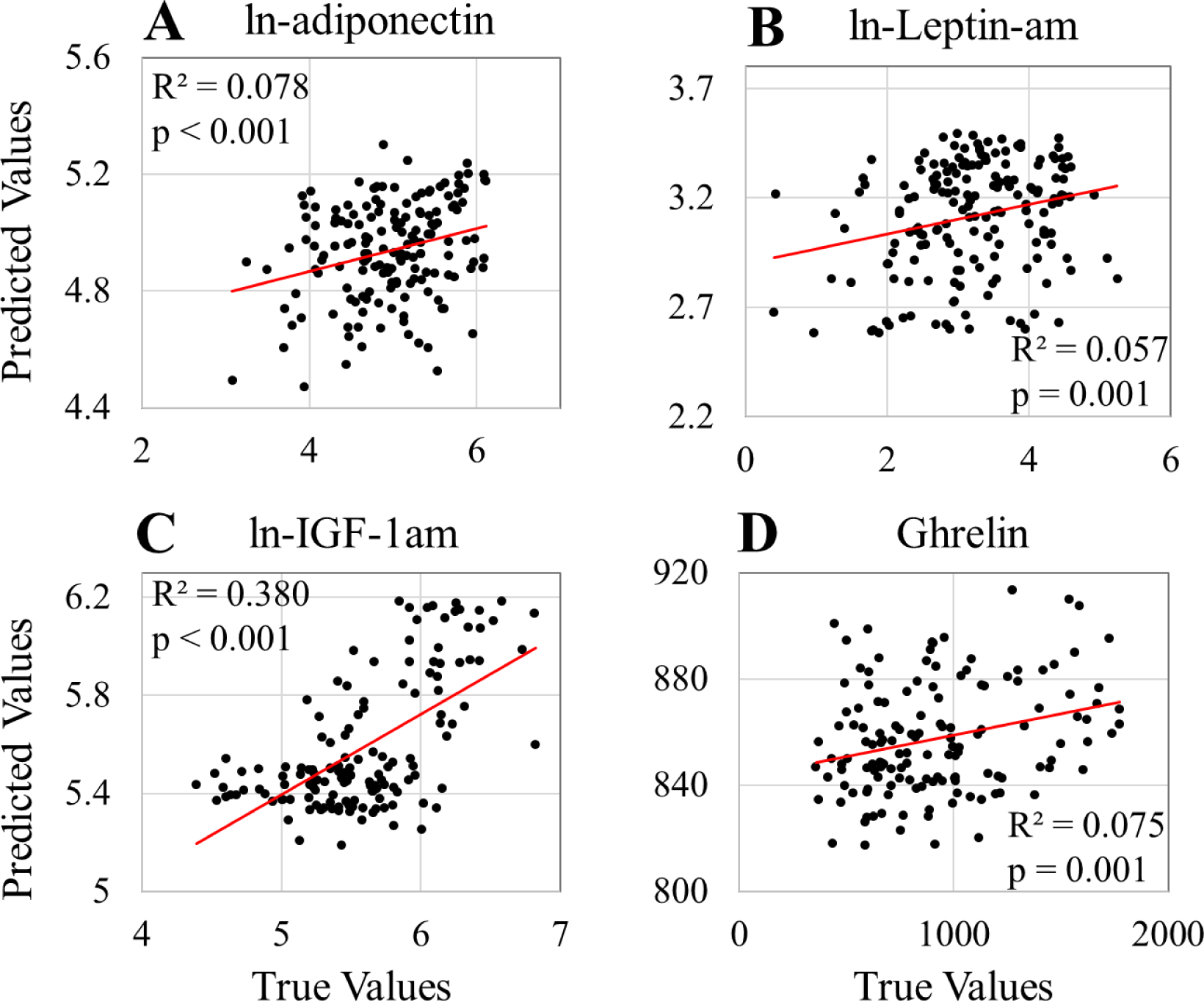
Regression plots for (A) ln-adiponectin, (B) ln-Leptin-am, (C) ln-IGF-1am, and (D) Ghrelin.

### Immune System Modeling

In predicting blood levels of ln-IL-6 using random forest regression, the model achieved an R² of 0.120 (p < 0.001), demonstrating the model’s ability to explain 12% of the variance in ln-IL-6 levels (Figure 5; Table 3). The predictive accuracy was characterized by a mean absolute error (MAE) of 0.567, mean squared error (MSE) of 0.501, root mean squared error (RMSE) of 0.708, and a mean absolute percentage error (MAPE) of 186.430%. For ln-TGF-alpha, regression analysis indicated an R² of 0.044 (p < 0.050), with an MAE of 0.514, MSE of 0.381, RMSE of 0.617, and MAPE of 110.426%. Similarly, predictions for ln-CRP levels yielded an R² of 0.084 (p < 0.001), an MAE of 1.014, MSE of 1.547, RMSE of 1.207, and MAPE of 218.306%. Regression models for blood ln-IL-1b and ln-IL-10 levels produced R² values of 0.006 (p = 0.436) and 0.007 (p = 0.338), respectively, indicating minimal explanatory power for the variance in these cytokine levels. The MAE, MSE, RMSE, and MAPE for ln-IL-1b were 0.714, 0.877, 0.936, and 127.201%, respectively. For ln-IL-10, the corresponding values were an MAE of 0.861, MSE of 1.187, RMSE of 1.089, and MAPE of 375.164%

**Figure 5.**
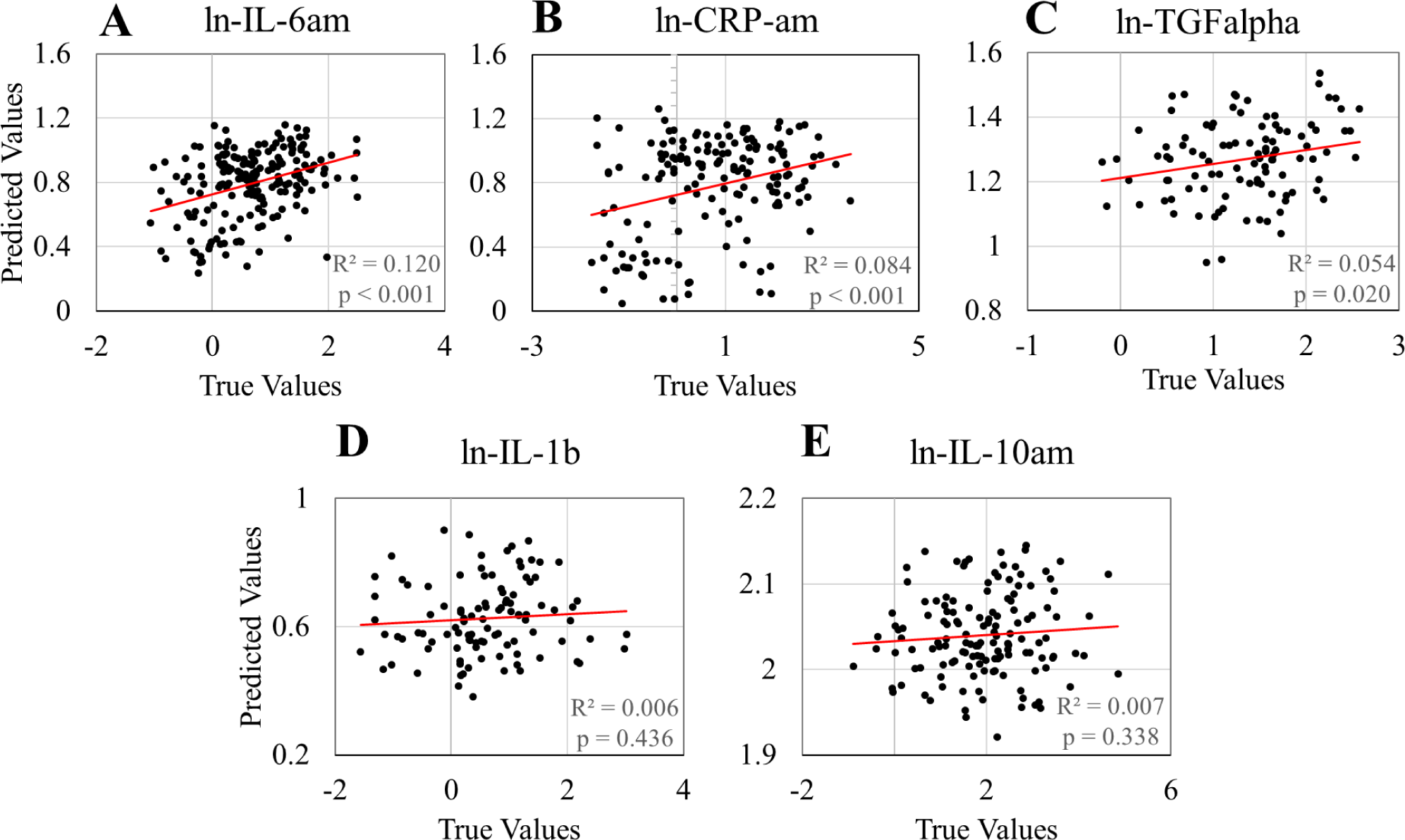
Regression plots for (A) ln-IL-6am, (B) ln-CRP-am, (C) ln-TGFalpha, (D) ln-IL-1b, and (E) ln-IL-10am.

### Sleep Apnea Modeling

Random forest regression analysis of apnea-hypopnea-index (AHI) values yielded an R² of 0.240 (p < 0.001), indicating the model accounted for 24% of the variance in AHI values (Figure 6; Table 3). The model exhibited a mean absolute error (MAE) of 0.924, mean squared error (MSE) of 1.357, root mean squared error (RMSE) of 1.165, and mean absolute percentage error (MAPE) of 129.173%.

**Figure 6.**
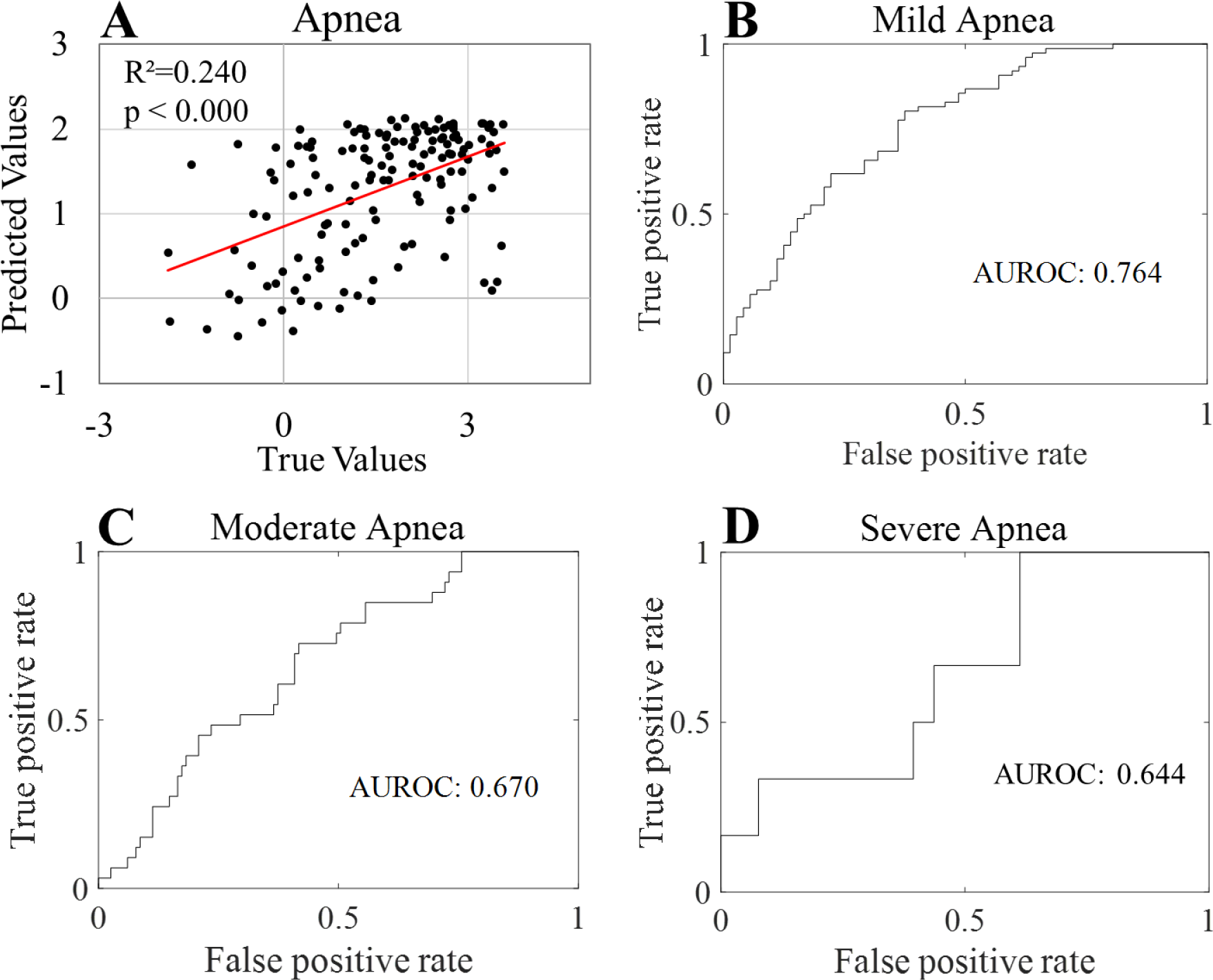
Regression and ROC plots for (A) all AHI values, (B) Mild apnea, (C) Moderate apnea, and (D) severe apnea.

Classification models for sleep apnea severity demonstrated varying levels of discriminative ability. For distinguishing individuals with mild or greater sleep apnea from those without apnea, the AUROC was 0.764, with an accuracy of 68.9% (Figure 6; Table 4). The precision for predicting mild apnea classification was 0.868 and that of predicting less than mild apnea was 0.5. Recall scores were 0.657 and 0.783 for mild apnea and no mild apnea, respectively, and their respective F1 scores were 0.742 and 0.610.

The model’s ability to classify individuals with moderate-or-greater severity of sleep apnea showed an AUROC of 0.670 and an accuracy of 56.1% (Figure 6; Table 4). The precision for predicting moderate apnea classification was 0.727 and that of predicting less than moderate apnea was 0.513. The recall scores for moderate apnea and no moderate apnea were calculated as 0.300 and 0.868, respectively, and their respective F1 scores were 0.425 and 0.645.

For severe sleep apnea, the AUROC was 0.644, and the model achieved a higher accuracy of 90.5% (although this value is influenced by the smaller sample size of severe apnea cases; Figure 6; Table 4). The precision for predicting severe apnea classification was 0.167, and that of predicting no severe apnea was 0.937. The recall scores for severe and no severe apnea classifications were 0.1 and 0.964, respectively, and their respective F1 scores were 0.125 and 0.95.

## Discussion

Our study demonstrates that features derived from oscillatory events in single-channel EEG are rich in biological information that informs our understanding of sleep-related physiological processes. We built our analyses on prior work that described morphological properties of slow waves and their temporal associations with coupled oscillatory events in the context of aging^30,51^, sex differences^37,38^, and endocrine function^40^. Leveraging these insights, we developed features that yield reproducible predictions in machine learning models for each of these processes. Our approach also demonstrated encouraging predictive results in several additional outcomes that are relevant to sleep and brain health monitoring, including cerebrovascular risk factors, inflammatory biomarkers, and sleep apnea.

In the field of machine learning applications that utilize sleep EEG metrics, prior studies predominantly relied on features such as spectral power composition and more basic morphological properties of slow waves and spindles^52–54^. Our analyses applied advanced methods to perform feature engineering from oscillatory events, enabling both subtyping of oscillatory events and more complex temporal associations of events that may increase the specificity and interpretability of predicted outcomes. Incorporating recent insights into how specific oscillatory events coordinate neural communication^24,25,44,55–57^, memory processing^58–60^, and synaptic regulation^61^, we created features that correlate with observable neural circuits^24,25,44,62^ and expanded specificity regarding the types of events that are analyzed. This method is highly advantageous in providing a basis for interpretable neuroanatomical localization of our findings. Further, the mapping of our feature extraction to definable brain communication events also facilitates the construction of digital biomarkers with more explainable biological foundations, contributing significantly to our understanding of the functional roles of oscillatory events in brain health monitoring.

Our oscillatory event-based feature set demonstrates successful predictive modeling within multiple previously described pathways connecting sleep with metabolic and hormonal systems. Sleep and circadian biology produce several fundamental regulatory components for insulin sensitivity and glucose homeostasis^39,63^, and indeed, experimental disruption of sleep has established a causal relationship between SWS and glucose dysregulation^11^. Preclinical models provide some mechanistic insights, including the involvement of hippocampal sharp wave ripples, an oscillatory event that occurs in conjunction with slow waves and sleep spindles, as an event that directly influences glucose homeostasis via hippocampal-hypothalamic pathways^64^. A sleep EEG-based analysis of slow wave and sleep spindle coupling also found a correlation between coupling metrics and insulin sensitivity^40^. Our analysis supports these findings, demonstrating success in predicting a component of variance in fasting glucose levels and glucose levels in OGTT. Our models also predicted levels of IGF-1, adiponectin, ghrelin, and leptin levels, which are closely related to the neurophysiological processes underpinning glucose homeostasis and insulin sensitivity^65–67^. Notably, our model for predicting insulin levels did not produce a significant regression, suggesting that our oscillatory event features may be more related to elements of glucose homeostasis beyond insulin secretion.

Predictive modeling was also successful in classification of health conditions including diabetes and hypertension, and our predictive modeling of BMI captured a component of biological variance as well. Our model performance in classification prediction for diagnosed diabetes suggests that oscillatory event features distinguish biological processes among individuals with chronic glucose control issues, in addition to the relationships identified with single-timepoint blood glucose levels. The prediction of hypertension diagnosis, although modest in performance, also indicates a potential for these features to assess the impact of blood pressure on brain health. Similarly, our predictive model for BMI indicates a likely relationship between oscillatory events and the health consequences of metabolic disorders associated with increased BMI. Notably, BMI has also been reported to impact the strength of electrocardiographic signals in sleep EEG^68^, and further studies may help assess whether similar technical aspects of EEG recording impact our feature extraction methods.

Our study also investigated the relationship between sleep apnea and oscillatory events. Prior reports have indicated that variations in slow wave activity and sleep spindles, such as changes in SWS duration and EEG power in specific frequency ranges, are associated with sleep apnea^69–72^. The models we developed showed modest predictive results with AHI, suggesting a correlation between oscillatory event features and the number of apneic and hypnotic events. Predicting binarized AHI severity grades was also modestly effective for mild, moderate, and severe thresholds of AHI, although with some limitations in the ability to model severe apnea due to lower availability of severe apnea recordings in the data set.

Sleep apnea is associated with a proinflammatory state^73^, which may in part explain some of the ability to predict cytokine levels in our models, although more direct relationships between cytokines and slow wave sleep are also likely to be important^14,15^. Given the associations between sleep apnea, BMI, and diabetes with proinflammatory states, it is plausible that our features are capturing biological processes related to these conditions and their association with immune biomarkers. Intriguingly, our models successfully predicted a component of the variance in IL-6, TNF-alpha, and CRP levels, but not IL-1b and IL-10 levels. This specificity in cytokine relationships to oscillatory event features warrants further investigation to examine reproducibility and possible mechanistic insights.

Sex differences in the morphological properties of slow waves have been described^37,38^, in addition to known differences in slow wave sleep architecture and age-related changes in slow wave sleep^38^. Consistent with these findings, our oscillatory event-based feature sets were effective in classification of sex and prediction of age. Future research is necessary to explore how environmental, genetic, and epigenetic differences may relate to predictive modeling of oscillatory events in differing demographic groups, highlighting the importance of developing unbiased and generalizable models in digital biomarker applications.

Our study focused on determining whether features derived from oscillatory events in single-channel EEG capture underlying biology, and our work should be considered an initial step in this feature engineering approach. Notably, our methods do not fully encompass all aspects of oscillatory event biology, such as more detailed subtyping of events and their temporal relationships, and the dynamic changes in oscillatory event across time series data. Additionally, while we used family ID to separate families into training or testing data to prevent model leakage, familial effects might still influence the models’ prediction capabilities, and the findings will need to be replicated independent of this potential confound. Future studies could further enhance generalizability and performance utilizing multiple cohorts during training and more sophisticated machine learning approaches, such as neural network-based predictive modeling.

Collectively, our results represent a pioneering effort in using domain knowledge of slow wave sleep oscillatory events for feature engineering and predictive modeling. These features, though relatively simple in their derivations, capture a significant amount of biological activity related to brain health in the machine learning predictions. Subsequent steps in developing digital biomarkers for monitoring brain health from single channel EEG may build on these features and enrich the feature sets for more specific properties of oscillatory events that represent dynamic properties of oscillatory events across cycles of sleep and through homeostatic mechanisms.

## References

1. Niedermeyer E, da Silva FL. Electroencephalography: basic principles, clinical applications, and related fields. Lippincot Williams & Wilkins; 2005.

2. Kourtis LC, Regele OB, Wright JM, Jones GB. Digital biomarkers for Alzheimer’s disease: the mobile/wearable devices opportunity. NPJ digital medicine. 2019;2(1):9.

3. Al-Qazzaz NK, Ali SHB, Ahmad SA, Chellappan K, Islam MS, Escudero J. Role of EEG as biomarker in the early detection and classification of dementia. The Scientific World Journal. 2014;2014

4. Cohen MX. *Analyzing neural time series data: theory and practice*. MIT press; 2014.

5. Brain A. Thalamocortical Oscillations in the Sleeping. SCIENCE. 1993;262:679.

6. Steriade M, McCormick DA, Sejnowski TJ. Thalamocortical oscillations in the sleeping and aroused brain. Science. 1993;262(5134):679–685.

7. Buzsaki G, Draguhn A. Neuronal oscillations in cortical networks. science. 2004;304(5679):1926–1929.

8. Léger D, Debellemaniere E, Rabat A, Bayon V, Benchenane K, Chennaoui M. Slow-wave sleep: from the cell to the clinic. Sleep medicine reviews. 2018;41:113–132.

9. Brodt S, Inostroza M, Niethard N, Born J. Sleep—A brain-state serving systems memory consolidation. Neuron. 2023;111(7):1050–1075.

10. Tononi G. Slow wave homeostasis and synaptic plasticity. Journal of Clinical Sleep Medicine. 2009;5(2 suppl):S16–S19.

11. Tasali E, Leproult R, Ehrmann DA, Van Cauter E. Slow-wave sleep and the risk of type 2 diabetes in humans. Proceedings of the National Academy of Sciences. 2008;105(3):1044–1049.

12. Born J, Fehm H. Hypothalamus-pituitary-adrenal activity during human sleep: a coordinating role for the limbic hippocampal system. Experimental and clinical endocrinology & diabetes. 1998;106(03):153–163.

13. Sassin J, Parker D, Mace J, Gotlin R, Johnson L, Rossman L. Human growth hormone release: relation to slow-wave sleep and sleep-waking cycles. Science. 1969;165(3892):513-515.

14. Besedovsky L, Lange T, Haack M. The sleep-immune crosstalk in health and disease. Physiological reviews. 2019;

15. Irwin MR, Opp MR. Sleep health: reciprocal regulation of sleep and innate immunity. Neuropsychopharmacology. 2017;42(1):129–155.

16. Pepin J-L, Borel A-L, Tamisier R, Baguet J-P, Levy P, Dauvilliers Y. Hypertension and sleep: overview of a tight relationship. Sleep medicine reviews. 2014;18(6):509–519.

17. Khalil M, Power N, Graham E, Deschênes SS, Schmitz N. The association between sleep and diabetes outcomes–A systematic review. Diabetes research and clinical practice. 2020;161:108035.

18. Nedeltcheva AV, Scheer FA. Metabolic effects of sleep disruption, links to obesity and diabetes. Current opinion in endocrinology, diabetes, and obesity. 2014;21(4):293.

19. 2020 Alzheimer’s disease facts and figures. Alzheimer’s & Dementia. 2020;16(3):391-460. doi:10.1002/alz.12068

20. Livingston G, Huntley J, Sommerlad A, et al. Dementia prevention, intervention, and care: 2020 report of the Lancet Commission. The Lancet. 2020;396(10248):413–446.

21. Irwin MR, Vitiello MV. Implications of sleep disturbance and inflammation for Alzheimer’s disease dementia. The Lancet Neurology. 2019;

22. Lee YF, Gerashchenko D, Timofeev I, Bacskai BJ, Kastanenka KV. Slow wave sleep is a promising intervention target for Alzheimer’s disease. Frontiers in neuroscience. 2020;14:705.

23. Bazhenov M, Timofeev I, Steriade M, Sejnowski T. Spiking-bursting activity in the thalamic reticular nucleus initiates sequences of spindle oscillations in thalamic networks. Journal of neurophysiology. 2000;84(2):1076–1087.

24. Jiang X, Gonzalez-Martinez J, Halgren E. Coordination of Human Hippocampal Sharpwave Ripples during NREM Sleep with Cortical Theta Bursts, Spindles, Downstates, and Upstates. Journal of Neuroscience. 2019;39(44):8744–8761.

25. Gonzalez CE, Mak-McCully RA, Rosen BQ, et al. Theta bursts precede, and spindles follow, cortical and thalamic downstates in human NREM sleep. Journal of Neuroscience. 2018;38(46):9989–10001.

26. Buzsáki G. Hippocampal sharp wave-ripple: A cognitive biomarker for episodic memory and planning. Hippocampus. 2015;25(10):1073–1188.

27. Chylinski D, Van Egroo M, Narbutas J, et al. Timely coupling of sleep spindles and slow waves linked to early amyloid-β burden and predicts memory decline. Elife. 2022;11:e78191.

28. Bouchard M, Lina J-M, Gaudreault P-O, et al. Sleeping at the switch. Elife. 2021;10:e64337.

29. Lafrenière A, Lina J-M, Hernandez J, Bouchard M, Gosselin N, Carrier J. Sleep slow waves’ negative-to-positive-phase transition: a marker of cognitive and apneic status in aging. Sleep. 2022;

30. Helfrich RF, Mander BA, Jagust WJ, Knight RT, Walker MP. Old brains come uncoupled in sleep: slow wave-spindle synchrony, brain atrophy, and forgeting. Neuron. 2018;97(1):221–230. e4.

31. Winer JR, Mander BA, Helfrich RF, et al. Sleep as a potential biomarker of tau and β-amyloid burden in the human brain. Journal of Neuroscience. 2019;39(32):6315–6324.

32. Pulver RL, Kronberg E, Medenblik LM, et al. Mapping sleep’s oscillatory events as a biomarker of Alzheimer’s disease. Alzheimers Dement. Aug 23 2023;doi:10.1002/alz.13420

33. Mander BA, Winer JR, Walker MP. Sleep and Human Aging. Neuron. 2017;94(1):19–36.

34. Mourtazaev M, Kemp B, Zwinderman A, Kamphuisen H. Age and gender affect different characteristics of slow waves in the sleep EEG. Sleep. 1995;18(7):557–564.

35. Dijk DJ, Beersma DG, Bloem GM. Sex differences in the sleep EEG of young adults: visual scoring and spectral analysis. Sleep. 1989;12(6):500–507.

36. Ehlers C, Kupfer D. Slow-wave sleep: do young adult men and women age differently? Journal of sleep research. 1997;6(3):211–215.

37. Carrier J, Viens I, Poirier G, et al. Sleep slow wave changes during the middle years of life. European Journal of Neuroscience. 2011;33(4):758–766.

38. Carrier J, Semba K, Deurveilher S, et al. Sex differences in age-related changes in the sleep-wake cycle. Frontiers in neuroendocrinology. 2017;47:66–85.

39. Broussard JL, Ehrmann DA, Van Cauter E, Tasali E, Brady MJ. Impaired insulin signaling in human adipocytes atier experimental sleep restriction: a randomized, crossover study. Annals of internal medicine. 2012;157(8):549–557.

40. Vallat R, Shah VD, Walker MP. Coordinated human sleeping brainwaves map peripheral body glucose homeostasis. Cell Reports Medicine. 2023;4(7)

41. Andrillon T, Nir Y, Staba RJ, et al. Sleep spindles in humans: insights from intracranial EEG and unit recordings. Journal of Neuroscience. 2011;31(49):17821–17834.

42. McConnell BV, Kronberg E, Medenblik LM, et al. The Rise and Fall of Slow Wave Tides: Vacillations in Coupled Slow Wave/Spindle Pairing Shiti the Composition of Slow Wave Activity in Accordance With Depth of Sleep. Original Research. Frontiers in Neuroscience. 2022-June-23 2022;16 doi:10.3389/fnins.2022.915934

43. Mak-McCully RA, Rolland M, Sargsyan A, et al. Coordination of cortical and thalamic activity during non-REM sleep in humans. Nature communications. 2017;8:15499.

44. Jiang X, Gonzalez-Martinez J, Halgren E. Posterior Hippocampal Spindle Ripples Co-occur with Neocortical Theta Bursts and Downstates-Upstates, and Phase-Lock with Parietal Spindles during NREM Sleep in Humans. Journal of Neuroscience. 2019;39(45):8949–8968.

45. Redline S, Tishler PV, Tosteson TD, et al. The familial aggregation of obstructive sleep apnea. American journal of respiratory and critical care medicine. 1995;151(3):682–687.

46. Zhang G-Q, Cui L, Mueller R, et al. The National Sleep Research Resource: towards a sleep data commons. Journal of the American Medical Informatics Association. 2018;25(10):1351–1358.

47. Patel SR, Zhu X, Storfer-Isser A, et al. Sleep duration and biomarkers of inflammation. Sleep. 2009;32(2):200–204.

48. Buxbaum SG, Elston RC, Tishler PV, Redline S. Genetics of the apnea hypopnea index in Caucasians and African Americans: I. Segregation analysis. Genetic Epidemiology: The Official Publication of the International Genetic Epidemiology Society. 2002;22(3):243–253.

49. Berry RB, Brooks R, Gamaldo CE, Harding SM, Marcus C, Vaughn B. The AASM manual for the scoring of sleep and associated events. *Rules, Terminology and Technical Specifications, Darien, Illinois*, American Academy of Sleep Medicine. 2012;

50. McConnell BV, Kronberg E, Teale PD, et al. The aging slow wave: a shitiing amalgam of distinct slow wave and spindle coupling subtypes define slow wave sleep across the human lifespan. Sleep. 2021;44(10):zsab125.

51. Ujma PP, Simor P, Steiger A, Dresler M, Bódizs R. Individual slow-wave morphology is a marker of aging. Neurobiology of aging. 2019;80:71–82.

52. Ye EM, Sun H, Krishnamurthy PV, et al. Dementia detection from brain activity during sleep. Sleep. 2023;46(3):zsac286.

53. Brink-Kjaer A, Leary EB, Sun H, et al. Age estimation from sleep studies using deep learning predicts life expectancy. NPJ digital medicine. 2022;5(1):103.

54. Sun H, Paixao L, Oliva JT, et al. Brain age from the electroencephalogram of sleep. Neurobiology of aging. 2019;74:112–120.

55. Maingret N, Girardeau G, Todorova R, Goutierre M, Zugaro M. Hippocampo-cortical coupling mediates memory consolidation during sleep. Nature neuroscience. 2016;19(7):959–964.

56. Andrade KC, Spoormaker VI, Dresler M, et al. Sleep spindles and hippocampal functional connectivity in human NREM sleep. Journal of Neuroscience. 2011;31(28):10331–10339.

57. Clemens Z, Mölle M, Erőss L, et al. Fine-tuned coupling between human parahippocampal ripples and sleep spindles. European Journal of Neuroscience. 2011;33(3):511–520.

58. Cairney SA, El Marj N, Staresina BP. Memory consolidation is linked to spindle-mediated information processing during sleep. Current Biology. 2018;28(6):948–954. e4.

59. Schreiner T, Petzka M, Staudigl T, Staresina BP. Endogenous memory reactivation during sleep in humans is clocked by slow oscillation-spindle complexes. Nature communications. 2021;12(1):1–10.

60. Latchoumane C-FV, Ngo H-VV, Born J, Shin H-S. Thalamic spindles promote memory formation during sleep through triple phase-locking of cortical, thalamic, and hippocampal rhythms. Neuron. 2017;95(2):424–435. e6.

61. Staresina BP, Niediek J, Borger V, Surges R, Mormann F. How coupled slow oscillations, spindles and ripples coordinate neuronal processing and communication during human sleep. Nature Neuroscience. 2023;26(8):1429–1437.

62. Verzhbinsky IA, Rubin DB, Kajfez S, et al. Co-occurring ripple oscillations facilitate neuronal interactions between cortical locations in humans. Proceedings of the National Academy of Sciences. 2024;121(1):e2312204121.

63. Broussard JL, Knud-Hansen BC, Grady S, et al. Influence of circadian phase and extended wakefulness on glucose levels during forced desynchrony. Sleep Health. 2023;

64. Tingley D, McClain K, Kaya E, Carpenter J, Buzsáki G. A metabolic function of the hippocampal sharp wave-ripple. Nature. 2021;597(7874):82–86.

65. Klok MD, Jakobsdotir S, Drent M. The role of leptin and ghrelin in the regulation of food intake and body weight in humans: a review. Obesity reviews. 2007;8(1):21–34.

66. Timper K, Brüning JC. Hypothalamic circuits regulating appetite and energy homeostasis: pathways to obesity. Disease models & mechanisms. 2017;10(6):679–689.

67. Muoio DM, Newgard CB. Molecular and metabolic mechanisms of insulin resistance and β-cell failure in type 2 diabetes. Nature reviews Molecular cell biology. 2008;9(3):193–205.

68. Purcell S, Manoach D, Demanuele C, et al. Characterizing sleep spindles in 11,630 individuals from the National Sleep Research Resource. Nature communications. 2017;8:15930.

69. Gaudreau H, Décary A, Sforza E, Petit D, Morisson F, Montplaisir J. Slow-wave activity in sleep apnea patients before and atier continuous positive airway pressure treatment: contribution to daytime sleepiness. Chest. 2001;119(6):1807–1813.

70. Himanen S-L, Virkkala J, Huupponen E, Hasan J. Spindle frequency remains slow in sleep apnea patientsthroughout the night. Sleep Medicine. 2003;4(3):229–234.

71. Ondze B, Espa F, Dauvilliers Y, Billiard M, Besset A. Sleep architecture, slow wave activity and sleep spindles in mild sleep disordered breathing. Clinical Neurophysiology. 2003;114(5):867–874.

72. Jones SG, Riedner BA, Smith RF, et al. Regional reductions in sleep electroencephalography power in obstructive sleep apnea: a high-density EEG study. Sleep. 2014;37(2):399–407.

73. Nadeem R, Molnar J, Madbouly EM, et al. Serum inflammatory markers in obstructive sleep apnea: a meta-analysis. Journal of Clinical Sleep Medicine. 2013;9(10):1003–1012.

